# D-mannose suppresses macrophage release of extracellular vesicles and ameliorates type 2 diabetes

**DOI:** 10.1101/2024.02.18.580856

**Authors:** Sha Zhang, Kai Zhang, Chen-Xi Zheng, Ying-Feng Gao, Guo-Rong Deng, Xu Zhang, Yuan Yuan, Ting Jia, Si-Yuan Tang, Guang-Xiang He, Zhen Gong, Cheng-Hu Hu, Bo Ma, Hong Zhang, Zhe Li, Yong-Chang Di-Wu, Yi-Han Liu, Liang Kong, Jing Ma, Yan Jin, Bing-Dong Sui

## Abstract

The monosaccharide D-mannose exists naturally in low abundance in human blood, while an increased plasma mannose level is associated with insulin resistance and the incidence of type 2 diabetes (T2D) in patients. However, whether and how D-mannose may regulate T2D development remains elusive. Here, we show that despite the altered mannose metabolism in T2D, drinking-water supplementation of supraphysiological D-mannose safely ameliorates T2D in genetically obese db/db mice. Interestingly, D-mannose therapy exerts limited effects on the gut microbiome and peripheral blood T cells, whereas D-mannose after administration is enriched in the liver and alleviates hepatic steatosis and insulin resistance. Mechanistically, D-mannose suppresses macrophage release of pathological extracellular vesicles (EVs) for improving hepatocyte function through metabolic control of CD36 expression. Collectively, these findings reveal D-mannose as an effective and potential T2D therapeutic, which add to the current knowledge of sugars regulating EV-based intercellular communication and inspire translational pharmaceutical strategies of T2D.

**Graphical abstract:** 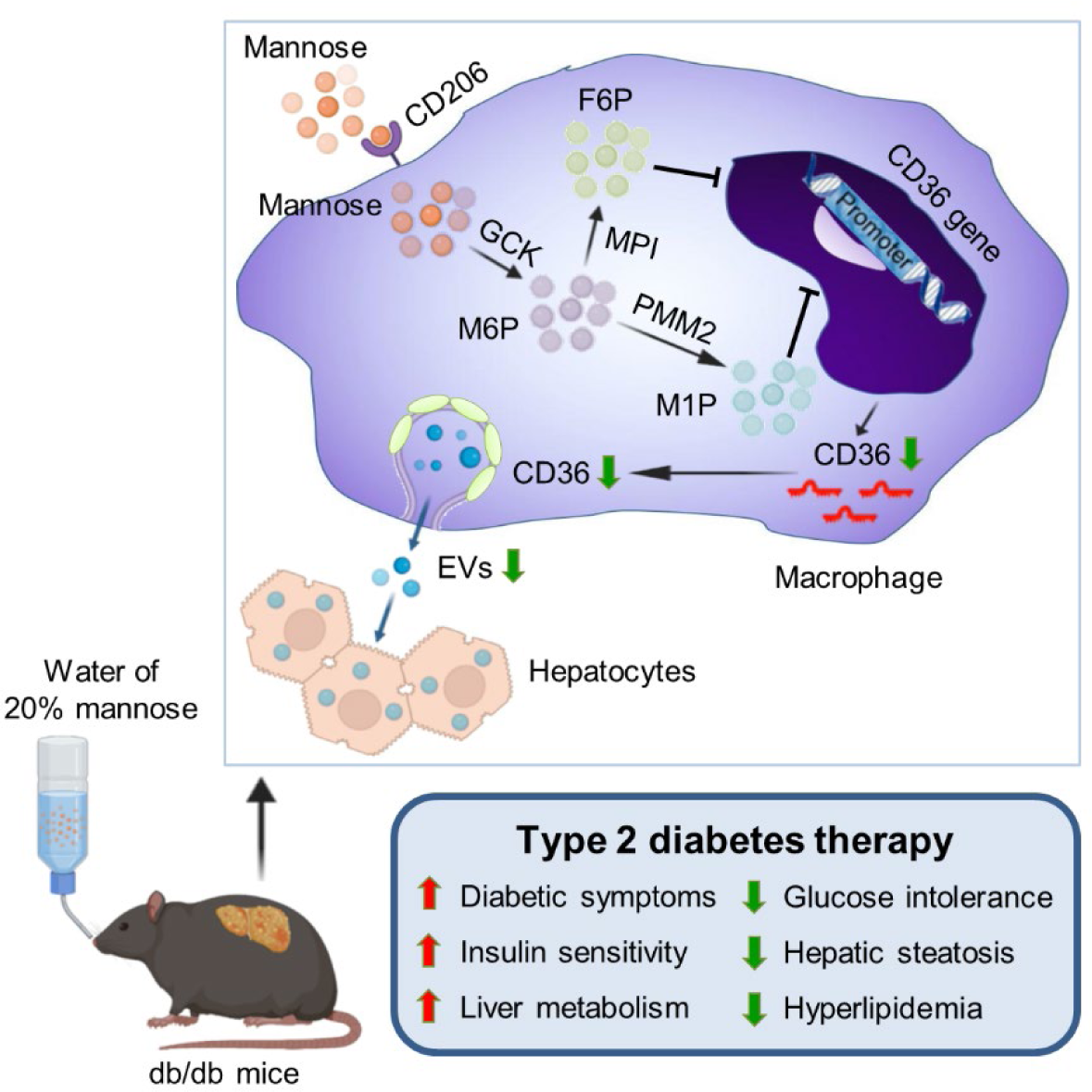

Drinking-water supplementation of D-mannose serves as an effective therapeutic of type 2 diabetes, which rescued hepatocyte steatosis through suppressing macrophage release of extracellular vesicles based on metabolic control of CD36 expression.

## 1. Introduction

Type 2 diabetes (T2D) is a chronic progressive metabolic disease with high and increasing global prevalence, which represents a major cause of morbidity and even mortality ^1,2^. T2D is characterized or associated with a wide spectrum of disease pathologies, including obesity, insulin resistance and macrophage-mediated chronic low-grade inflammation ^1,3^. Particularly, pro-inflammatory activation of tissue macrophages in the obese condition is known to release multiple cytokines, such as tumor necrosis factor-alpha (TNF-α) and galectin-3, as well as extracellular vesicles (EVs), membranous nanoparticles for intercellular communication, to impair insulin sensitivity and induce metabolic alterations (*e.g.*, hepatic steatosis) in target organs ^4–6^. However, current anti-inflammatory therapies have limited effects on ameliorating insulin resistance in T2D patients ^7–9^. Furthermore, although the role of endogenous EVs in human health and disease is emergingly being revealed, clinically available pharmaceuticals for controlling pathological EV release is still lacking ^10,11^. Therefore, there remains an unmet need to unravel feasible pharmacological targets of restraining macrophage-based paracrine crosstalk in T2D and develop therapeutic approaches accordingly.

Integrated analyses by combining cell-specific genome-scale metabolic, transcriptional regulatory and protein-protein interaction networks in human have identified increased levels of plasma mannose in obese subjects and discovered significant correlations between high circulating mannose concentrations with insulin resistance and the incidence of T2D in large prospective cohorts ^12,13^. D-mannose, a natural C-2 epimer of glucose, is a monosaccharide found in plants and fruits and exists in human blood at a concentration less than one-fiftieth of that of glucose, which contributes to protein glycosylation and represents an inefficient cellular energy source ^14–16^. Importantly, D-mannose administration orally *via* drinking water at supraphysiological levels has been proved effective as clinical therapeutics for patients with the mannose phosphate isomerase (MPI)-congenital disorder of glycosylation (MPI-CDG) and recurrent urinary tract infection (UTI) ^17,18^, which has also been reported useful to treat T lymphocyte- and macrophage-associated immunopathologies and improve glucose and lipid metabolism in mice ^19–22^. Therefore, D-mannose might exert beneficial effects on T2D despite the increased plasma level, but whether and how D-mannose regulates T2D development remains elusive. Notably, the release of EVs has been emergingly revealed to be regulated by sugars and glycosylation ^23^. Further investigations on the potential effects of D-mannose modulating EV release under pathophysiological condition may provide additional interesting mechanisms supporting its translational promise.

In this study, we aim to investigate that whether and how D-mannose may regulate the T2D development. Through a series of experiments in the genetically obese, leptin receptor-deficient db/db mice, we show that despite the altered mannose metabolism in T2D, drinking-water supplementation of D-mannose at the supraphysiological level safely ameliorates T2D. Interestingly, D-mannose therapy exerts limited effects on the gut microbiome and peripheral blood T lymphocytes, whereas D-mannose after administration is enriched in the liver and ameliorates hepatic steatosis and insulin resistance. Mechanistically, D-mannose inhibits macrophage release of EVs for improving hepatocyte function through metabolic control of CD36 expression. Collectively, these findings reveal D-mannose as an effective and potential T2D therapeutic, which add to the current knowledge of sugars regulating EV-mediated intercellular communication and shed light on translational pharmaceutical strategies of T2D.

## 2. Materials & Methods

### 2.1 Mice

BKS.Cg-*Dock7^m^* +/+ *Lepr^db^*/J mice (strain NO. 000642) were purchased from the Jackson Laboratory, USA ^24^. Non-obese and non-diabetic heterozygotes from the colony (denoted as db/m) were used as the control and for breeding of the obese and diabetic *Lepr^db^*/+ *Lepr^db^* (denoted as db/db) homozygotes. Male mice were used from 5-week old to 13-week old, which were housed in pathogen-free conditions, maintained on a standard 12-h light-dark cycle, and received normal chow diet and water *ad libitum*. All animal experiments were performed in compliance with the relevant laws and ethical regulations, following the Guidelines of Intramural Animal Use and Care Committees of The Fourth Military Medical University, approved by the Ethics Committee of The Fourth Military Medical University, and following the ARRIVE guidelines.

### 2.2 Cell lines

The RAW 264.7 mouse macrophage cell line was obtained from the American Type Culture Collection (TIB-71; ATCC, USA). Cells were cultured in Dulbecco’s Modified Eagle Medium with 1 g/L D-glucose (low-glucose DMEM; Invitrogen, USA) supplemented with 10% fetal bovine serum (FBS; ExCell Bio, China), 2 mM L-glutamine (Invitrogen, USA) and 1% penicillin/streptomycin (Invitrogen, USA) and incubated at 37°C under 5% CO2.

### 2.3 Primary cell cultures

For culture of primary macrophages from the bone marrow (BMDMs) ^25^, long bones of mice were harvested and bone marrow cavities were flushed with phosphate-buffered saline (PBS; Invitrogen, USA), which was then passed through a cell strainer and subjected to red blood cell lysis (Solarbio, China). Freshly isolated cells were cultured in low-glucose DMEM (Invitrogen, USA) supplemented with 10% FBS (ExCell Bio, China), 2 mM L-glutamine (Invitrogen, USA), 1% penicillin/streptomycin (Invitrogen, USA) and 20 ng/ml recombinant mouse macrophage-colony stimulating factor (M-CSF; PeproTech, USA). After induction for 7 days, mature BMDMs were collected and used for collection of mEVs.

Isolation of primary mouse hepatocytes was performed by perfusion *via* the portal vein ^5^. Briefly, mouse liver was perfused *via* catheterization of the portal vein using a 24G needle catheter (BD, USA) and a mini-pump machine (Thermo Fisher Scientific, USA) under general anesthesia. The liver was perfused firstly with 10 ml Hank’s balanced salt solution (HBSS) (Invitrogen, USA) to remove blood followed by 20 ml HBSS supplemented with 1 mM ethylene glycolbis(aminoethylether)-tetra-acetic acid (EGTA) (Sigma-Aldrich, USA) to remove the endogenous calcium. Then the liver was perfused with 20 ml HBSS supplemented with 5 mM calcium chloride (CaCl2) (Sigma-Aldrich, USA) and 40 μg/ml liberase TM (Sigma-Aldrich, USA) for digestion. All the solutions were kept at 37°C in a water bath. After digestion, the liver was dissected and washed in ice-chilled HBSS, and cells were teased out into DMEM with 4.5 g/L D-glucose (high-glucose DMEM; Invitrogen, USA) supplemented with 10% FBS (Sigma-Aldrich, USA) and 1% penicillin/streptomycin (Invitrogen, USA). Hepatocytes were then prepared by centrifugation at 50 g for 5 min at 4°C, filtered through 70 μm nylon strainers, purified by a 49% Percoll solution (Sigma-Aldrich, USA), and resuspended in William’s E Medium (WEM) with GlutaMAX™ (Invitrogen, USA) containing 10% FBS (Sigma-Aldrich, USA), 10 nM dexamethasone (Invitrogen, USA) and 1% penicillin/streptomycin (Invitrogen, USA). Hepatocytes were then seeded onto collagen-coated plates (Corning, USA) or coverslips (Electron Microscopy Sciences, USA) and incubated at 37°C in a humidified atmosphere of 5% CO2 overnight for attachment.

### 2.4 Chemical treatments

As reported ^19^, drinking-water supplementation of D-mannose was performed by dissolving 20 g D-mannose (Shanghai Yuanye Bio-Technology, China) in 100 mL distilled water (20% or 0.2 g/mL, equal to 1.1 mol/L), and unsupplemented control water was given as the control. *In vitro* treatment of D-mannose was used at the concentration of 25 mM for 48 h based on our preliminary dose-effect tests on inhibiting EV release (data not shown) and published papers on immunomodulation of T cells and macrophages ^19,20^. After internalization, D-mannose is phosphorylated by hexokinase to produce M6P, which undergo two major metabolic fates: a minor fraction (∼5%) is isomerized to M1P by PMM2 to be used in glycosylation pathways; the large majority (∼95%) is converted to F6P by MPI to be catabolized into glycolysis ^16^. Therefore, to investigate the D-mannose metabolic effect, Tunicamycin (MedChemExpress, China) was used to block protein N-glycosylation ^26^ at 100 ng/mL for 48 h, and MLS0315771 (MedChemExpress, China) was applied to suppress MPI ^27^ at 5 mM for 48 h, the dose and duration were selected based on our preliminary dose-effect tests on macrophage viability (data not shown). Furthermore, to test the effects of D-mannose metabolites, M6P (Aladdin, China) was used at 25 mM (equal to the D-mannose concentration) for 48 h, M1P (Aladdin, China) was used at 1.25 mM (5% of the D-mannose dose) for 48 h, and F6P (Yingxinbio, China) was used at 23.75 mM (95% of the D-mannose dose) for 48 h. PA (Kunchuang Biotechnology, China) was added at 500 μM for 24 h based on our preliminary dose-effect tests on macrophage viability (data not shown) and previous research on regulating RAW 264.7 cells and EV release ^28,29^.

### 2.5 CD36 overexpression

The overexpression of CD36 was performed using the lentivirus-based vector by Hanbio, China, with the vector being used as the negative control. After transfection at a multiplicity of infection (MOI) of 50, all RAW264.7 cells were treated with puromycin (Solarbio, China) at a concentration of 10 μg/mL for 10 days. The lentivirus transfection efficacy was validated by fluorescent imaging, qRT-PCR and Western blot analysis.

### 2.6 Isolation, labeling and treatment of mEVs

EVs were isolated from cultured macrophages based on our established protocol ^30^. Briefly, cells were cultured in complete medium containing EV-depleted FBS for 48 h. EV-depleted FBS was obtained by ultracentrifugation at 100,000 g for 18 h. The culture supernatant was collected and subsequently centrifuged at 800 g for 10 min. The supernatant was further collected and centrifuged at 16,500 g for 30 min at 4°C to obtain mEVs, which were then washed with filtered PBS. mEV pellets collected from each six-well were photographed, and quantification of mEVs was performed using the BCA method (Beyotime, China) for protein amounts. The lipophilic dye PKH67 (Sigma-Aldrich, USA) was used to label mEVs according to the manufacturer’s instructions and according to our previous report ^25^. In specific, after PKH67 staining for 5 min, mEV suspension in PBS was added by an equal volume of EV-depleted FBS and incubated for 1 min to allow binding of excess PKH67 dye. mEVs were then collected *via* centrifugation and washed with PBS to get rid of unbound PKH67. The supernatant was used as the negative control, and the mEV pellets were resuspended for usage. For *in vitro* treatment, mEVs were dissolved in PBS and added to the culture medium at a protein concentration of 20 μg/ml, with the dose being determined by preliminary tests on recipient hepatocyte viability (data not shown) and our previous experience ^25^. For *in vivo* treatment, mEVs were dissolved in PBS and were infused *via* the caudal vein into recipient mice at 200 μg on the basis of protein measurement ^25^ every 5 days during the experimental period.

### 2.7 Gross analysis and blood glucose quantification

Mice were recorded for body weight and water and food intake every 3 days. Cage bedding were photographed every 5 days. The concentrations of non-fasting random blood glucose were measured every 3 days throughout the experiments, and the concentrations of fasting blood glucose were measured after fasting for 6 h ^24,31^. Blood glucose was quantified using an ACCU-CKEK glucometer (Roche, Germany) following tail vein-puncture of whole blood sampling. The concentrations of HbA1c were determined using the A1CNow Self Check system (Sinocare, China). All mice were euthanasia at the end of experiments, photographed for gross view images, collected for organs and quantified for organ weights. Fat mass and lean mass were determined using the minispec LF90 Whole Body Composition Analyzer (Bruker, Germany) before euthanasia.

### 2.8 IPGTT and IPITT assays

IPGTT and IPITT were performed based on our previous study with minor modifications ^25^. For IPGTT, mice were fasted for 20 h and intraperitoneally injected with D-glucose (Oubokai, China) at 1.5 g/kg. For IPITT, mice were fasted for 6 h and intraperitoneally injected with recombinant human insulin (Novo Nordisk, Denmark) at 2 IU/kg. Blood glucose levels were measured at 0, 15, 30, 60, 90 and 120 min after D-glucose or insulin administration.

### 2.9 Lipid content measurement

For hepatic lysate examination, at sacrifice, liver tissues were dissected at approximately 100 mg and homogenized in 9-fold volume of ethanol on ice. The lysates were centrifuged at 2,500 g for 10 min at 4°C, and the supernatant was collected. For serum examination, at sacrifice, whole peripheral blood was extracted from the mouse retro-orbital venous plexus, and serum was isolated by centrifugation at 3,000 g for 15 min at 4°C. TG, TC and FFA levels were measured by the commercial kits according to the manufacturer’s instructions (Nanjing Jiancheng Biology Engineering Institute, China) ^25^.

### 2.10 ELISA

Plasma was isolated from extracted whole peripheral blood by adding heparin solution (STEMCELL Technologies, USA) followed by centrifugation at 1,300 g for 15 min at 4°C. Concentrations of TNF-α and IL-10 were determined using commercial kits (Fankewei, China) according to the manufacturer’s instructions.

### 2.11 Peripheral blood T-cell analysis

Whole peripheral blood was extracted from the mouse retro-orbital venous plexus with anti- coagulation, and cells were isolated by centrifugation at 500 g for 5 min at 4°C followed by being treated with a red blood lysis buffer (Beckman Coulter, USA). After washing with PBS, PBMNCs were collected by centrifugation at 500 g for 5 min at 4°C. PBMNCs were then stained with fluorescence-conjugated antibodies for CD3, CD4 and CD8 (all from BioLegend, USA) at 1:100 for 30 min at 4°C in dark, washed and examined by the ACEA NovoCyte flow cytometer (Agilent, USA).

### 2.12 Flow cytometric EV analysis

According to our published protocol ^32,33^, collected plasma was further centrifuged at 2,500 g for 10 min at 4°C to remove the platelets after being diluted with the same volume of PBS. The supernatant was then centrifuged at 16,500 g for 30 min at 4°C to pellet EVs. The pellet was resuspended and washed with 0.2 μm-filtered PBS, stained with the FITC-conjugated F4/80 antibody (BioLegend, USA) at 1:50 for 30 min at 4°C in dark, washed and examined by the ACEA NovoCyte flow cytometer (Agilent, USA). Collected mEVs in PBS were stained with fluorescence-conjugated antibodies for F4/80 and CD11b (both from BioLegend, USA) at 1:50 for 30 min at 4°C in dark, washed and also examined by the ACEA NovoCyte flow cytometer (Agilent, USA).

### 2.13 HPLC analysis

D-mannose concentrations in serum and liver were examined by HPLC analysis. For serum, samples were added with distilled water at 1:2 (v/v), mixed by ultrasound treatment for 30 min on ice, and centrifuged at 12,000 g for 10 min at 4°C. The supernatant was collected. For hepatic lysates, liver tissues were dissected at approximately 100 mg and homogenized in 500 μL distilled water on ice. The lysates were centrifuged at 12,000 g for 10 min at 4°C, and the supernatant was collected. HPLC examination was performed on 10 μL samples at a flow rate of 1.0 mL/min with HP-Amino columns (Sepax Technologies, USA).

### 2.14 Biodistribution analysis

Cy5.5-labeled mannose (Qiyue Biology, China) was oral gavaged at 10 mg, and mice were euthanatized after 24 h. The organs were harvested and imaged using the IVIS Lumina XRMS Series 2 instrument (PerkinElmer, USA) to assess the biodistribution of mannose, and the fluorescence intensity was quantified using the Living Image software (PerkinElmer, USA)^25^. The liver was then subjected to standard IF staining for cellular uptake analysis, as stated below.

### 2.15 Histological analysis

At sacrifice, multiple organs were isolated and fixed overnight with 4% paraformaldehyde (PFA) (Saint-bio, China). Samples were dehydrated and embedded in paraffin, and 5 μm serial sections were prepared (RM2125; Leica, Germany). Sections then underwent H&E staining using a commercial kit (Beyotime, China). Hepatic steatosis was graded blindly based on the NAFLD activity score, which was performed by assessing the percentage of hepatocytes containing lipid droplets (S0: less than 5%; S1: 5-33%; S2: 34-66%; S3: greater than 66%) ^34^.

### 2.16 ORO and IF staining

At sacrifice, liver tissues were rapidly isolated, fixed overnight in 4% PFA, cryoprotected with 30% (w/v) sucrose (Solarbio, China), and embedded in optimal cutting temperature (OCT) compound (Sakura Finetek, USA). The specimens were snap-frozen and sectioned into 10 μm sagittal sections (CM1950; Leica, Germany). For ORO staining ^25^, liver sections were immersed in 3 mg/mL ORO working solution (Aladdin, China) for 5 min and rinsed with distilled water. Sections were counterstained with the hematoxylin solution (Beyotime, China), mounted and photographed with a microscope (M205FA; Leica, Germany). Percentages of lipid droplet area were quantified using ImageJ 1.47 software (National Institute of Health, USA). For F-actin staining of hepatocyte borders, as previously stated ^35^, sections were probed with phalloidin conjugated to AlexaFluor-568 (Invitrogen, USA) according to the manufacturer’s instructions, and counterstained with 4’,6-diamidino-2-phenylindole (DAPI) (Abcam, UK). For IF staining of the protein markers, sections were treated with 0.3% Triton X-100 (Solarbio, China) diluted in PBS for 20 min at room temperature, blocked with 10% goat serum (Boster, China) for 1 h at room temperature, and stained with a rabbit anti-mouse TNF-α primary antibody (Novus Biologicals, USA), a rat anti-mouse CD206 primary antibody (Bio-Rad, USA), or a rat anti-mouse F4/80 primary antibody (Abcam, UK) overnight at 4°C at a concentration of 1:100. After washing with PBS, sections were then stained with a FITC-AffiniPure Goat Anti-Rabbit IgG secondary antibody (Yeasen, China) or a FITC-AffiniPure Goat Anti-Rat IgG secondary antibody (Yeasen, China) for 1 h at room temperature at a concentration of 1:200, and counterstained with DAPI (Abcam, UK). Fluorescent images were obtained using a confocal laser scanning microscope (Olympus, Japan).

### 2.17 Hepatic insulin signaling examination

Hepatic insulin signaling was evaluated by measuring insulin-stimulated AKT and AMPKα phosphorylation ^25^. Briefly, mice were fasted for 8 h and then injected intraperitoneally with recombinant human insulin (Novo Nordisk, Denmark) at 1 IU/kg. Mice were sacrificed at 15 min after injection, and liver tissues were collected and snap frozen with liquid nitrogen. The phosphorylation of AKT and AMPKα was determined by Western blot analysis, as stated below.

### 2.18 RNA extraction and qRT-PCR analysis

Total RNA was isolated from freshly isolated liver tissues after homogenization under liquid nitrogen or from cultured RAW 264.7 using the commercial RNA isolation kits (Foergene, China) according to the manufacturer’s instructions. The cDNA was synthesized using a commercial a PrimeScript^TM^ RT Reagent Kit (Takara, Japan). Then, qRT-PCR was performed with a SYBR Premix Ex Taq II Kit (Takara, Japan) by a Real-Time System (CFX96; Bio-Rad, USA). Quantification was performed by using β-actin as the internal control and calculating the relative expression level of each gene with the 2^-ΔΔCT^ method, as previously described ^25^. Primers used in this study were listed in Table S3.

### 2.19 Western blot analysis

Western blot analysis was performed according to our previous study ^25^. Whole lysates of liver tissues, RAW 264.7 cells or EVs were prepared using the RIPA Lysis Buffer (Beyotime, China). Proteins were extracted and the protein concentration was quantified using the BCA method (Beyotime, China). Equal amounts of protein samples were loaded onto SDS-PAGE gels and transferred to polyvinylidene fluoride (PVDF) membranes (Millipore, USA) which were blocked with 5% bovine serum albumin (BSA) (Gemini Bio, USA) in TBS for 2 h at room temperature. Then, the membranes were incubated overnight at 4°C with the following primary antibodies: anti-AMPKα (Cell Signaling Technology, USA; diluted at 1:1000), anti-p- AMPKα (Thr172) (Cell Signaling Technology, USA; diluted at 1:1000), anti-p-AKT (Ser473) (Cell Signaling Technology, USA; diluted at 1:1000), anti-AKT (Cell Signaling Technology, USA; diluted at 1:1000), anti-CD36 (Abmart, China; diluted at 1:1000), anti-CD9 (HuaBio, China; diluted at 1:1000), anti-CD63 (Thermo Fisher Scientific, USA; diluted at 1:1000), anti- CD81 (GeneTex, USA; diluted at 1:1000), anti-Cav-1 (Santa Cruz Biotechnology, USA; diluted at 1:1000), anti-Mitofilin (Abcam, UK; diluted at 1:1000), anti-Golgin84/GOLGA5 (Novus Biologicals, USA; diluted at 1:1000), anti-GAPDH (Proteintech, China; diluted at 1:1000), anti-Vinculin (Abmart, China; diluted at 1:1000) and anti-β-actin antibodies (Proteintech, China; diluted at 1:2500). After washing with TBS containing 0.1% Tween-20, the membranes were incubated with horseradish peroxidase (HRP)-conjugated secondary antibodies (Signalway, China) for 1 h at room temperature. The protein bands were visualized using an enhanced chemiluminescence kit (Amersham Biosciences, USA) and detected by a gel imaging system (4600; Tanon, China).

### 2.20 NTA assay

NTA was performed by a ZetaView instrument (Particle Metrix, Germany). Resuspended mEVs were diluted 50-fold in filtered PBS to achieve a final concentration of 3.2×10^10^ particle/mL. The capture length was 60 s with camera level set to 14 and detection threshold set to 3. The image of filtered PBS was taken to verify that the diluent had no particle in it. A total of 1498 frames were captured and analyzed ^25^. The ZetaView software (Particle Metrix, Germany) was used for capturing and data analysis.

### 2.21 TEM analysis

mEV pellet was resuspended in 2% PFA, and 20 μl mEVs were deposited on 200-mesh formvar-coated copper grids and dried at room temperature for 5 min. After removing excess suspension using filter paper, the mEVs were negatively stained with uranyl acetate (Sigma-Aldrich, USA) at room temperature for 2 min, washed with distilled water and dried. Imaging was performed under a FEI Tecnai G2 Spirit Biotwin TEM (Thermo Fisher Scientific, USA) operating at 100 kV, with a PHURONA camera (EMSIS, Germany) and RADIUS 2.0 software (EMSIS, Germany) ^25^.

### 2.22 Metabolic assays of hepatocytes

For the glucose uptake assay ^36^, hepatocytes after overnight attachment were changed to DMEM medium with no D-glucose (Invitrogen, USA) for overnight incubation. A glucose uptake commercial kit based on the fluorescent glucose analog 2- (N- (7-nitrobenz-2-oxa- 1,3-diazol-4-yl)amino)-2-deoxyglucose (2-NBDG) (Cayman Chemical, USA) was used to determine the glucose uptake rate. Glucose output of hepatocytes was also evaluated after overnight incubation in DMEM medium with no D-glucose (Invitrogen, USA) ^37^. Hepatocytes were then treated with 2 mM sodium pyruvate (Sigma-Aldrich, USA) and 20 mM sodium lactate (Sigma-Aldrich, USA) for 4 h, and supernatant of the culture medium was collected and examined using a glucose determination commercial kit (Sigma-Aldrich, USA). For fatty acid uptake ^38^, a commercial kit (Abnova, USA) was used based on a fluorescent fatty acid substrate, which was added to cultured hepatocytes in BSA-free medium with the fluorescence intensity being read immediately using the microplate reader (Synergy H1; Bio- Tek, USA) with the Gen5 software (Bio-Tek, USA) at 485/515 nm kinetically for 60 min, at an interval of 1 min. Lipid output of hepatocytes was also evaluated in BSA-free medium after adding 0.5 mM sodium acetate (Sigma-Aldrich, USA) for 2 h ^39^. Supernatant of the culture medium was collected and examined using the commercial kit for TG (Nanjing Jiancheng Biology Engineering Institute, China).

### 2.23 16S rRNA sequencing

Based on the previous study ^21^, mouse fecal samples were collected with aseptic techniques, and total genomic DNA was extracted from the feces. PCR amplification of the V3-V4 region of bacterial 16S rRNA was performed using 341F and 805R primers. Quality control was performed on raw data to obtain high-quality clean data for subsequent analyses. Alpha diversity was calculated using the QIIME2 microbiome bioinformatics platform. Beta diversity was also calculated using QIIME2, and the graphics were plotted using R packages (v3.5.2).

The sequence alignment was performed using Blast, and representative sequences were annotated using the SILVA database. All amplicon sequence variants (ASVs) identified in the 16S rRNA sequencing were listed in Table S1.

### 2.24 RNA sequencing analysis

Macrophages were treated with or without D-mannose at 25 mM for 48 h and then washed with PBS for 3 times. Total RNA was isolated using Trizol (Invitrogen, USA) according to the manual instruction. RNA sequencing libraries were generated with an insert size ranging from 100 to 500 bp, and sequenced using the BGISEQ-500 platform (Bioprofile, China) ^25^. In the Linux environment, FastQC (version 0.12.1) was used for quality control filtering. STAR (version 2.7.11a) was used to align clean reads to the reference genome. Gene abundance was represented as fragments per kilobase million (FPKM). In the R environment, DESeq2 package (version 1.4.5) was used for differentially expressed gene (DEG) (log2Fold change > 0.3 and adjusted p-value < 0.05) analysis. Pheatmap package (version 1.0.12) was used to generate the heatmap for the DEGs. The details of all the identified genes were listed in Table S2.

### 2.25 Network pharmacology

The target collection for D-mannose was obtained from online databases, including Swiss Target Prediction (PMID: 31106366), Similarity Ensemble Approach (PMID: 17287757), TargetNET (PMID: 27167132), BindingDatabase (PMID: 17145705), Therapeutic Target Database (PMID: 37713619), DrugBank (PMID: 18048412), STITCH (PMID: 18084021), ChEMMBL (PMID: 21948594), and Pharmmapper (PMID: 20430828). Targets of T2D were identified using databases, such as MalaCards (PMID: 23584832), OMIM (PMID: 15608251), PharmGKB (PMID: 34387941), Therapeutic Target Database (PMID: 37713619), DigSeE (PMID: 23761452), DrugBank (PMID: 18048412), DisGenet (PMID: 31680165), and GeneCards (PMID: 20689021). The target genes of D-mannose in the interaction with T2D were determined by the intersection of D-mannose and T2D target gene sets. The Venn diagram was generated to visualize the overlapping genes using an online tool (https://bioinfogp.cnb.csic.es/tools/venny/index.html). GO enrichment analysis was performed using the enrichment network tool. The top 20 GO enrichment items were listed according to the q-value (adjusted p-value), and the results were presented in a scatter plot using the Appyters network application.

### 2.26 Quantification and Statistical Analysis

Data were presented as mean ± standard deviation (SD) or as box (25th, 50th, and 75th percentiles) and whisker (range) plots of at least three independent experiments or three biological replicates. Data were analyzed by two-tailed unpaired Student’s *t* test for two-group comparisons, one-way analysis of variation (ANOVA) followed by the Turkey’s post-hoc test for multiple comparisons, or Kruskal-Wallis test for non-parametric comparisons using the Prism 5.01 software (GraphPad, USA). *P* values of less than 0.05 were considered statistically significant.

***Note:*** *All antibody, Chemicals, Critical Commercial Assays, Medium, Cell Lines, Animals, Oligonucleotides, Recombinant DNA, Software and Algorithms information used in the paper can be found in Table S4*.

## 3. Results

### 3.1 Altered mannose metabolism is associated with T2D in genetically obese db/db mice

To begin, we selected the genetically obese db/db mice as the representative T2D mouse model ^24^, and male mice were used to exclude the potential side effects caused by estrogen. The db/db mice, expectedly, developed increasingly higher body weight from 5 weeks to 13 weeks of age (*i.e.*, the experimental period) compared to their age- and sex-matched db/m control (Figure 1A and B). Furthermore, during the experimental period, db/db mice exhibited high random blood glucose levels over the diabetic criteria of 16.8 mmol/L (about 300 mg/dL), with also high fasting blood glucose levels over the diabetic criteria of 11.1 mmol/L (about 200 mg/dL) (Figure 1C and D). Moreover, percentages of glycated hemoglobin A1C (Hb1Ac) in blood, which indicates long-term glucose status, were detected over the diabetic criteria of 6.5% in db/db mice (Figure 1E). Notably, db/db mice resembled clinical diabetic symptoms of polyuria, polydipsia and polyphagia, showing wetter bedding and higher amount of water and food intake than db/m mice (Figure 1F-H). Importantly, db/db mice demonstrated reduced tolerance to intraperitoneal glucose injection and decreased insulin sensitivity, as evidenced by intraperitoneal glucose and insulin tolerance tests (IPGTT and IPITT, respectively) (Figure 1I-L).

**Figure 1.**
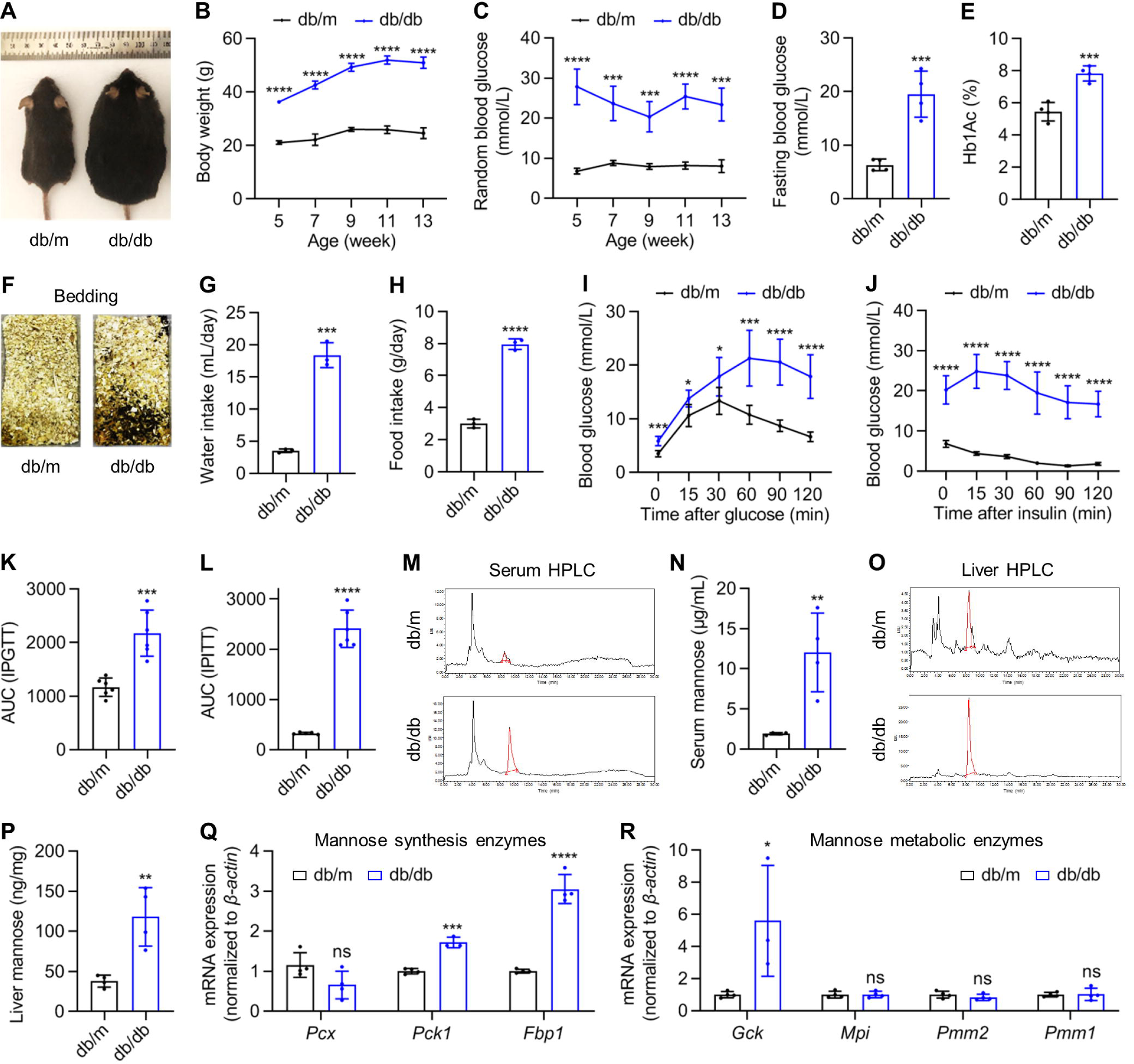
Altered mannose metabolism is associated with type 2 diabetes (T2D) in genetically obese db/db mice. (A) The gross view image of db/m and db/db male mice at 13-week old. (B) Body weight changes of db/m and db/db male mice. n=4. (C) Random blood glucose levels sampled from the tail vein. n=4. (D) Blood glucose levels after fasting for 6 h. n=4. (E) Blood hemoglobin A1C (Hb1Ac) levels. n=4. (F) Cage bedding after 5 days of 4 mice in each cage. (G) Average water intake of a single mouse per day. n=3. (H) Average food intake of a single mouse per day. n=3. (I) The intraperitoneal glucose tolerance test (IPGTT) after 20-h fasting and 1 g/kg glucose injection. n=5-6. (J) The intraperitoneal insulin tolerance test (IPITT) after 6-h fasting and 2 IU/kg insulin injection. n=5-6. (K) Area under curve (AUC) analysis of IPGTT. n=5-6. (L) AUC analysis of IPITT. n=5-6. (M) High performance liquid chromatography (HPLC) analysis of serum samples with red peaks indicating mannose. (N) Serum mannose levels quantified by HPLC. n=4. (O) HPLC analysis of liver contents with red peaks indicating mannose. (P) Liver mannose levels quantified by HPLC. n=4. (Q) Quantitative real-time polymerase chain reaction (qRT-PCR) analysis of mannose synthesis enzyme expression in the liver. n=3-4. (R) qRT-PCR analysis of mannose metabolic enzyme expression in the liver. n=3-4. Mean ± SD. *, *P* < 0.05; **, *P* < 0.01; ***, *P* < 0.001; ****, *P* < 0.0001; ns, *P* > 0.05. Two-tailed unpaired Student’s *t* test.

For mannose levels, we applied high performance liquid chromatography (HPLC) to examine serum samples (Figure 1M). Results revealed that the average levels of mannose in serum of db/db mice were 6-fold higher than db/m mice, reaching 12 μg/mL (beyond 60 μM) with statistical significance (Figure 1N). HPLC examination of mannose contents in the liver, the major organ of physiological mannose synthesis and metabolic consumption ^12,40^, also demonstrated over 2-fold increase in db/db mice compared to db/m mice (Figure 1O and P). Accordingly, quantitative real-time polymerase chain reaction (qRT-PCR) analysis of mRNA expression of mannose synthesis enzymes in the liver, namely the critical genes for gluconeogenesis ^41^, *Pyruvate carboxylase* (*Pcx*), *Phosphoenolpyruvate carboxykinase 1* (*Pck1*) and *Fructose bisphosphatase 1* (*Fbp1*), revealed upregulation of *Pck1* and *Fbp1* in db/db mice (Figure 1Q). Interestingly, mRNA expression levels of mannose metabolic enzymes in the liver ^16^, namely *Glucokinase* (*Gck*), *Mpi*, *Phosphomannomutase 2* (*Pmm2*) and *Phosphomannomutase 1* (*Pmm1*), showed upregulated or paralleled levels in db/db mice (Figure 1R). These results indicated maintained capability of metabolizing mannose in T2D, despite the non-specific acceleration of mannose production attributed to promoted gluconeogenesis. Taken together, these findings suggest that altered mannose metabolism is associated with T2D in db/db mice.

### 3.2 Drinking-water supplementation of D-mannose safely ameliorates T2D in db/db mice

The maintained capability of metabolizing mannose in T2D and the previously documented multiple beneficial effects of D-mannose enlightened us to investigate whether D-mannose administration would help alleviate T2D in db/db mice. Pioneer studies have established that supraphysiological concentrations of mannose (20% or 0.2 g/mL, equal to 1.1 mol/L) can be safely achieved by drinking-water supplementation ^19^. Therefore, we adopted this method to treat db/db mice during the 8-week experimental period (Figure 2A). Intriguingly, we found that drinking-water supplementation of D-mannose did not significantly affect the body weight gain (Figure 2B and C) or the random blood glucose levels (Figure 2D) of db/db mice, but it indeed reduced the fasting blood glucose levels and rescued the blood Hb1Ac percentages of db/db mice (Figure 2E and F), indicating efficacy related to stimulation and in the long-term. Furthermore, oral administration of D-mannose ameliorated the diabetic symptoms in db/db mice, showing controlled urine output and suppressed water and food intake (Figure 2G-I). Importantly, D-mannose administration improved the glucose tolerance condition and promoted insulin sensitivity of db/db mice, as evidenced by IPGTT and IPITT, suggesting the therapeutic effects (Figure 2J-M). In addition, histological analysis across multiple organs identified limited influence of D-mannose on the heart, lung, kidney and spleen morphology (Figure S1A-H), despite reduced organ weights in db/db mice, with also limited effects on the fat mass or lean mass of db/db mice (Figure S1I and J). Taken together, these findings indicate that drinking-water supplementation of D-mannose safely ameliorates T2D in db/db mice.

**Figure 2.**
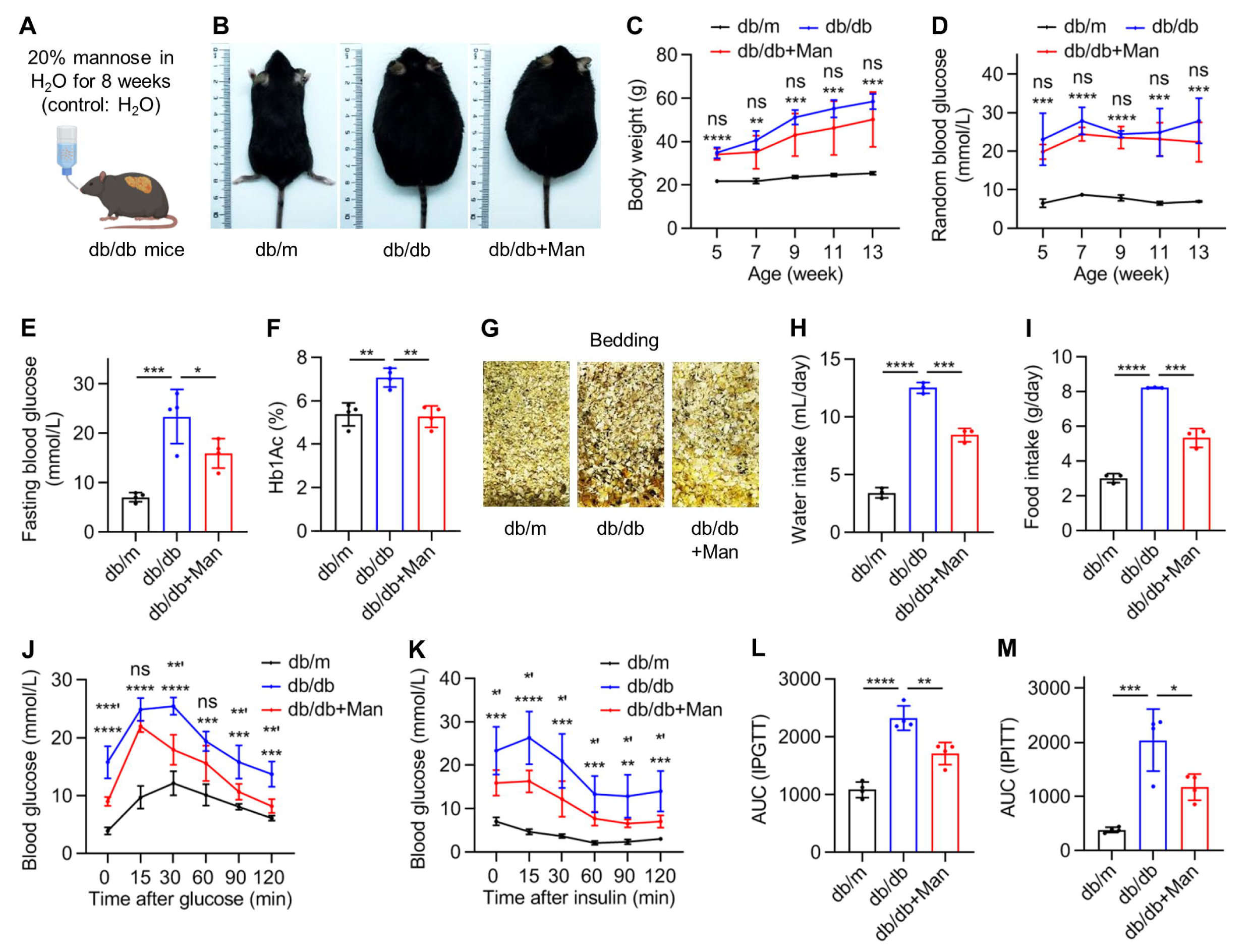
Drinking-water supplementation of D-mannose safely ameliorates type 2 diabetes (T2D) in db/db mice (Figure S1-related). (A) The diagram showing drinking-water supplementation of D-mannose (Man). (B) Gross view images of mice at 13-week old. (C) Body weight changes of mice. n=4-5. **, *** or ****, db/db compared to db/m. ns, db/db+Man compared to db/db. (D) Random blood glucose levels sampled from the tail vein. n=4. *** or ****, db/db compared to db/m. ns, db/db+Man compared to db/db. (E) Blood glucose levels after fasting for 6 h. n=4. (F) Blood hemoglobin A1C (Hb1Ac) levels. n=4. (G) Cage bedding after 5 days of 4 mice in each cage. (H) Average water intake of a single mouse per day. n=3. (I) Average food intake of a single mouse per day. n=3. (J) The intraperitoneal glucose tolerance test (IPGTT) after 20-h fasting and 1 g/kg glucose injection. n=4. *** or ****, db/db compared to db/m. **’, ***’ or ns, db/db+Man compared to db/db. (K) The intraperitoneal insulin tolerance test (IPITT) after 6-h fasting and 2 IU/kg insulin injection. n=4. **, *** or ****, db/db compared to db/m. *’, db/db+Man compared to db/db. (L) Area under curve (AUC) analysis of IPGTT. n=4. (M) AUC analysis of IPITT. n=4. Mean ± SD. * and *’, *P* < 0.05; ** and **’, *P* < 0.01; *** and ***’, *P* < 0.001; ****, *P* < 0.0001; ns, *P* > 0.05. One-way ANOVA with Turkey’s post-hoc test.

### 3.3 D-mannose administration exerts limited effects on the gut microbiome and peripheral blood T lymphocytes

Next, we investigated how D-mannose may ameliorate T2D in db/db mice. Previous studies have reported that mannose administration in water regulates gut microbiome and prevents high-fat diet (HFD)-induced obesity, and that drinking-water supplementation of D-mannose suppresses T lymphocytes-based immunopathology for autoimmune type 1 diabetes (T1D) therapy ^19,21^. With this knowledge, we first performed biodistribution analysis of Cy5.5-labeled fluorescent mannose after oral administration in db/db mice. Data showed that exogenous mannose mainly distributed in the liver after 24 h, suggesting successful entrance into circulation (Figure 3A). Notably, the bowel and the kidney were also fluorescently labeled, which corresponded to absorption and excretion of mannose respectively through the intestine and *via* the urine (Figure 3A). Accordingly, we examined that whether the gut microbiome in db/db mice was affected by mannose administration. 16S rRNA sequencing (Table S1) and related indexes of α diversity showed limited influence of D-mannose on the gut microbiome of db/db mice (Figure 3B-E), which were further supported by the principal coordinates analysis (PCoA) and the nonmetric multidimensional scaling (NMDS) analyses of β diversity (Figure 3F). Moreover, the relative abundance of microbiome compositions with quantifications at phylum and genus levels of *Firmicutes* over *Bacteroidetes* ratio, a relevant index of gut dysbiosis in obese individuals ^42^, revealed no significant effects of D-mannose administration (Figure 3G-J). Considering that D-mannose entered into the blood after intestinal absorption, we then examined that whether peripheral blood T cells were affected. Flow cytometric analysis demonstrated that neither the total CD3^+^ T cell percentages among the peripheral blood mononucleated cells (PBMNCs) (Figure 3K) nor the CD4^+^/CD8^+^ T cell ratios (Figure 3L and M) was influenced by D-mannose. Taken together, these data suggest that D-mannose administration exerts limited effects on the gut microbiome and peripheral blood T lymphocytes.

**Figure 3.**
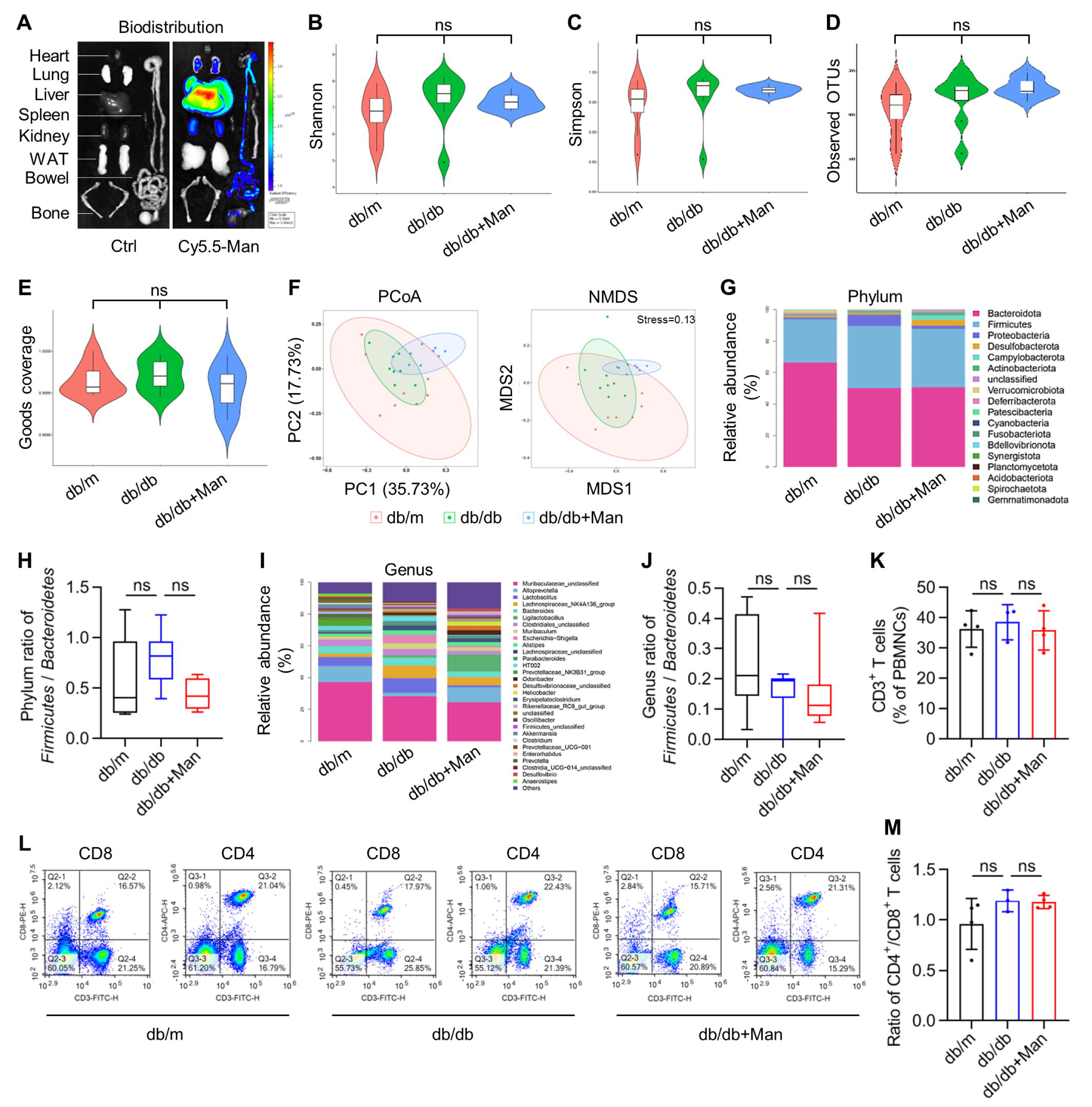
D-mannose administration exerts limited effects on the gut microbiome and peripheral blood T lymphocytes. (A) Biodistribution of Cy5.5-labeled fluorescent mannose (Man) after oral administration for 24 h. WAT, white adipose tissue. (B) The violin plot showing the shannon index of α diversity in 16S rRNA sequencing of the gut microbiome. n=8. (C) The violin plot showing the simpson index of α diversity. n=8. (D) The violin plot showing the observed operational taxonomic units (OTUs) index of α diversity. n=8. (E) The violin plot showing the goods coverage index of α diversity. n=8. (F) Principal coordinates analysis (PCoA) and nonmetric multidimensional scaling (NMDS) analyses of β diversity. n=8. (G) The stacked bar chart showing the relative abundance of phyla. (H) Quantification of ratio of *Firmiccutes* over *Bacteroidetes* at the phylum level. n=6-8. (I) The stacked bar chart showing the relative abundance of genera. (J) Quantification of ratio of *Firmiccutes* over *Bacteroidetes* at the genus level. n=6-8. (K) CD3^+^ T cell percentages in peripheral blood mononucleated cells (PBMNCs) analyzed by flow cytometry. n=3-4. (L) Flow cytometric analysis of CD8^+^ and CD4^+^ percentages in peripheral blood CD3^+^ T cells. (M) Quantification of ratio of CD4^+^ T cells over CD8^+^ T cells in the peripheral blood. n=3-4. ns, *P* > 0.05. Box (25th, 50th, and 75th percentiles) and whisker (range) and Kruskal-Wallis test (B-E, H and J), or mean ± SD and One-way ANOVA with Turkey’s post-hoc test (K and M).

### 3.4 D-mannose therapy alleviates hepatic steatosis and insulin resistance

The liver is a major metabolic organ which pathologically undergoes metabolic dysfunction, develops fatty liver disease, reveals insulin resistance and contributes to T2D ^43^. As liver is also the primary organ for circulatory mannose consumption (Figure 3A) ^40^, we next evaluated whether D-mannose improved hepatic conditions in treating T2D. Gross analysis detected that the db/db liver had steatosis appearance, while D-mannose administration benefited the liver status (Figure 4A). D-mannose amelioration of the fatty liver in db/db mice was further confirmed at the histological level by hematoxylin and eosin (H&E) staining (Figure 3B), as well as oil red O (ORO) staining (Figure 3C), which revealed decreased lipid droplet area in the D-mannose-treated liver. Accordingly, D-mannose reduced the liver / body weight ratio (Figure 4D) and alleviated hepatic steatosis with statistical significance in db/db mice (Figure 4E and F). Moreover, the elevated contents of triglyceride (TG), total cholesterol (TC) and free fatty acids (FFA) in the db/db liver were rescued by D-mannose therapy (Figure 4G-I), which were further correlated with recovered hyperlipidemia (Figure 4J-L). We have also examined the hepatic insulin resistance by performing Western blot analysis after insulin injection. Data demonstrated that phosphorylation levels of AKT and adenosine 5’- monophosphate (AMP)-activated protein kinase alpha (AMPKα) were decreased in the db/db liver, which suggested impaired insulin sensitivity and were rescued by D-mannose therapy (Figure 4M). Notably, D-mannose administration also restored expression related to hepatic glucose output and lipid metabolism, including glucose-6-phosphatase (encoded by *Glucose- 6-phosphatase catalytic subunit 1*, *G6pc1*), carbohydrate response element binding protein (ChREBP, encoded by *MLX interacting protein like*, *Mlxipl*), peroxisome proliferator-activated receptor gamma (PPARγ, encoded by *Pparg*) and PPARγ coactivator-1alpha (PGC-1α, encoded by *Ppargc1*). Taken together, these results suggest that D-mannose therapy alleviates hepatic steatosis and insulin resistance.

**Figure 4.**
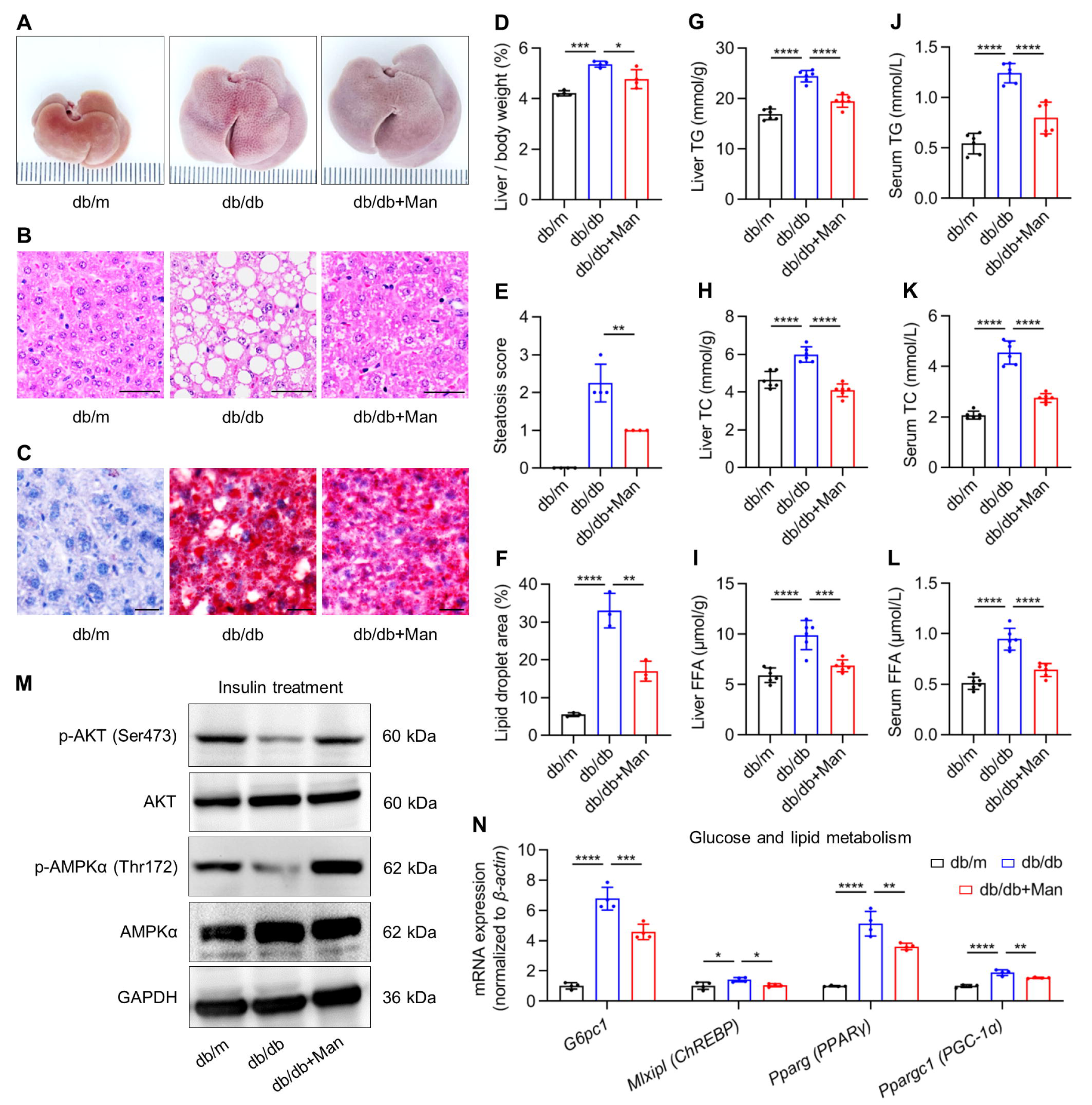
D-mannose therapy alleviates hepatic steatosis and insulin resistance. (A) Gross view images of the liver. Man, mannose. (B) Hematoxylin and eosin (H&E) staining images showing the liver histology. Scale bars, 100 μm. (C) Oil red O staining images showing lipid droplets in the liver. Scale bars, 25 μm. (D) Ratio of liver weight over body weight. n=4. (E) Quantification of hepatic steatosis in H&E staining images. n=4. (F) Quantification of lipid droplet area percentages in oil red O staining images. n=3. (G) Liver triglyceride (TG) contents analyzed by chemical tests. n=6. (H) Liver total cholesterol (TC) contents analyzed by chemical tests. n=6. (I) Liver free fatty acid (FFA) contents analyzed by chemical tests. n=6. (J) Serum TG contents analyzed by chemical tests. n=6. (K) Serum TC contents analyzed by chemical tests. n=6. (L) Serum FFA contents analyzed by chemical tests. n=6. (M) Western blot analysis of phosphorylation levels of AKT and adenosine 5’- monophosphate (AMP)-activated protein kinase alpha (AMPKα) in the liver after 1 IU/kg insulin treatment for 15 min. (N) Quantitative real-time polymerase chain reaction (qRT-PCR) analysis of glucose and lipid metabolic gene expression in the liver. n=4. Mean ± SD. *, *P* < 0.05; **, *P* < 0.01; ***, *P* < 0.001; ****, *P* < 0.0001. One-way ANOVA with Turkey’s post-hoc test.

### 3.5 D-mannose inhibits macrophage release of EVs for improving hepatocyte function]

Next, we deciphered how D-mannose may improve hepatic steatosis. The mannose receptor (also termed CD206) is predominantly expressed on macrophages, among other cells, and modulates their polarization and inflammatory response ^44^. Accordingly, we have performed immunofluorescent (IF) staining and discovered that F4/80-marked liver macrophages were the main cells internalizing Cy5.5-labeled mannose (Figure 5A). We have also adopted the network pharmacology method to predict molecular targets of D-mannose in treating T2D, which showed 138 overlapped genes between D-mannose and T2D (Figure 5B). Of these potential targets, Gene Oncology (GO) enrichment analysis revealed that most of the Top 20-enriched terms were related to EVs (Figure 5C), which was notable and surprising. Thus, we investigated that whether liver macrophages were regulated by D-mannose therapy and that whether macrophage release of EVs was involved. Further IF staining of macrophage activation/polarization markers in the liver, TNF-α (pro-inflammatory) and CD206 *per se* (anti-inflammatory) (Figure 5D), as well as enzyme-linked immunosorbent assay (ELISA) of plasma TNF-α and interleukin-10 (IL-10, anti-inflammatory) concentrations (Figure 5E and F), exhibited that D-mannose suppressed the pro-inflammatory reaction of liver macrophages without affecting the anti-inflammatory response. Particularly, db/db mice showed a notable characteristic of increased F4/80^+^ macrophage-derived EV population in circulation, which was restored by D-mannose therapy (Figure 5G). These results suggest that paracrine effects of macrophages, especially EV release, are involved in D-mannose therapy of T2D.

**Figure 5.**
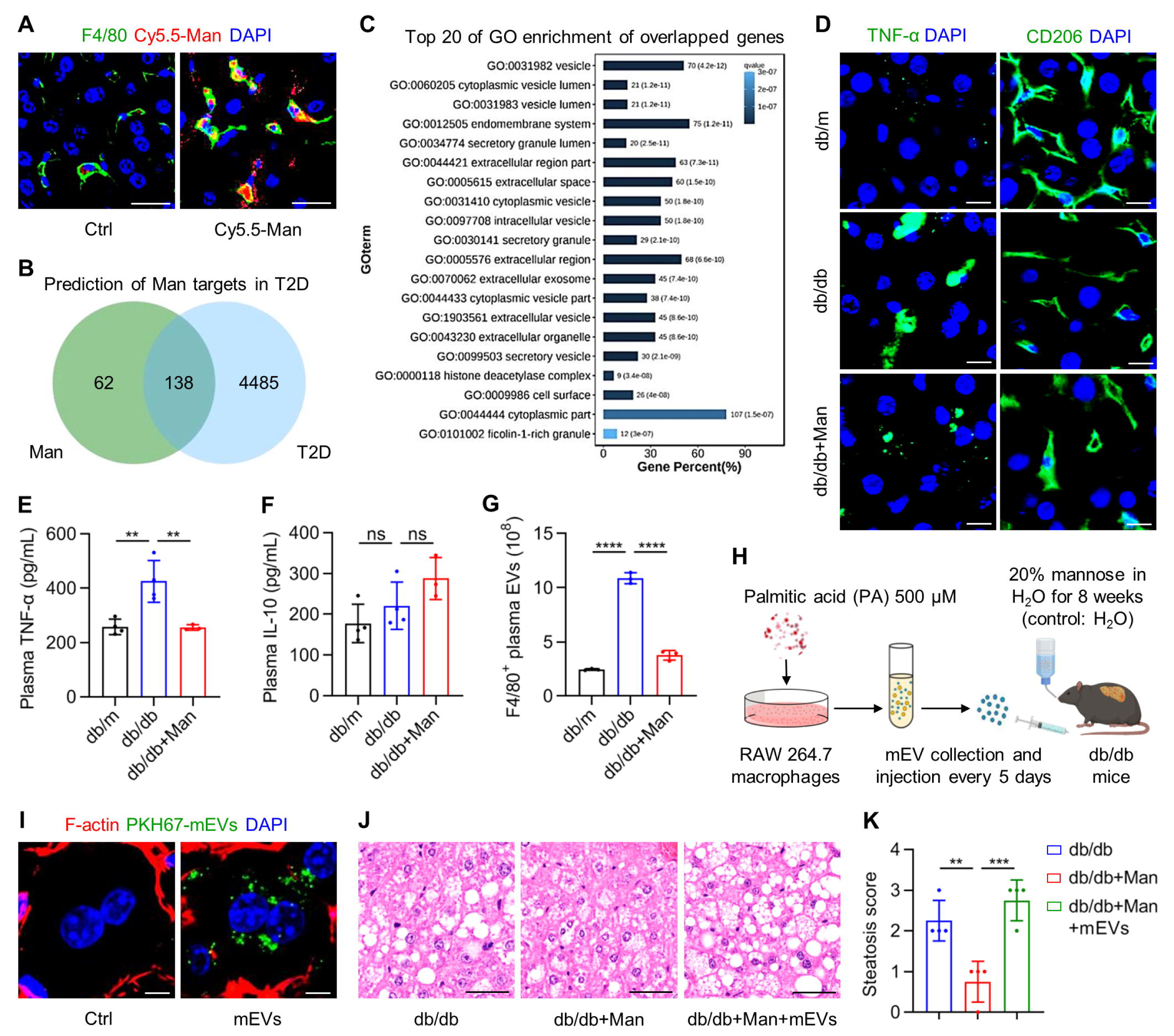
D-mannose inhibits macrophage release of extracellular vesicles (EVs) for improving hepatocyte function (Figure S2-related). (A) Fluorescent images showing the internalization of Cy5.5-labeled mannose (Man, red) by F4/80-marked macrophages (green) in the liver, counteracted by DAPI (blue). Scale bars, 50 μm. (B) Prediction of potential mannose targets in type 2 diabetes (T2D) by network pharmacology. (C) Top 20 terms of Gene oncology (GO) enrichment analysis of overlapped genes of mannose and T2D. (D) Fluorescent images showing tumor necrosis factor-alpha (TNF-α) or CD206 (green) positive area in the liver, counteracted by DAPI (blue). Scale bars, 25 μm. (E) Enzyme-linked immunosorbent assay (ELISA) of plasma TNF-α levels. n=3-4. (F) ELISA of plasma interleukin-10 (IL-10) levels. n=3-4. (G) Quantification of F4/80^+^ macrophage-produced EVs in the plasma by nanoparticle tracking analysis (NTA) combined with flow cytometric analysis. n=3. (H) The diagram showing the experimental procedure of macrophage-derived EV (mEV) collection and injection. PA, palmitic acid. (I) Fluorescent images showing the uptake of PKH67-labeled mEVs (green) by F-actin-labeled hepatocytes (red) in the liver, counteracted by DAPI (blue). Scale bars, 5 μm. (J) Hematoxylin and eosin (H&E) staining images showing the liver histology. Scale bars, 100 μm. (K) Quantification of hepatic steatosis in H&E staining images. n=4. Mean ± SD. **, *P* < 0.01; ***, *P* < 0.001; ****, *P* < 0.0001; ns, *P* > 0.05. One- way ANOVA with Turkey’s post-hoc test.

To confirm whether D-mannose regulation of macrophage release of EVs was indeed critical to its therapeutic efficacy, we cultured the RAW264.7 mouse macrophage cell line, treated them with palmitic acid (PA), a saturated fatty acid to mimick the T2D environment *in vitro*, collected macrophage-derived EVs (mEVs) by differential centrifugation, and injected mEVs intermittently back into D-mannose-administered db/db mice (Figure 5H). The isolated mEVs demonstrated featured size distribution ranging from 50-500 nm peaked at 100-200 nm (Figure S2A), a typical cup-shaped morphology (Figure S2B), expression of macrophage surface markers (Figure S2C) and representative EV proteins (Figure S2D) ^32^. Fluorescent tracing of PKH67-labeled mEVs in the recipient liver identified that almost all infused mEVs were detected within the F-actin-marked hepatocyte cell border, indicating uptake of mEVs by hepatocytes (Figure 5I). Accordingly, treatment of cultured hepatocytes with mEVs (Figure S2E) demonstrated that mEVs from the db/db macrophages inhibited glucose and fatty acid uptake while promoting glucose and lipid output of hepatocytes (Figure S2F-I), underlying the development of T2D. Expectedly, replenishment of PA-preconditioned mEVs in D-mannose-treated db/db mice blocked therapeutic effects of D-mannose on hepatic steatosis, leaving pathological lipid droplet deposition despite D-mannose administration (Figure 5J and K). Taken together, these findings suggest that D-mannose inhibits macrophage release of EVs for improving hepatocyte function.

### 3.6 D-mannose metabolism suppresses CD36 expression in macrophages to control EV release

Finally, we dissected how D-mannose may regulate macrophage release of EVs. By treating RAW 264.7 mouse macrophages with PA and D-mannose (Figure 6A), we confirmed that the number of mEVs released, rather than their protein content, was promoted by PA and was restored by D-mannose (Figure 6B-E). To explore the potential molecule(s) mediating effects of D-mannose, we performed next-generation RNA-sequencing analysis on macrophages, and the transcriptome data suggested multiple genes modulated, of which CD36, a recently reported regulator of EV release and fatty acid uptake, was involved (Figure 6F) ^45^. qRT-PCR analysis of CD36 expression in cultured macrophages confirmed its upregulation after PA treatment, which was suppressed by D-mannose (Figure 6G). Protein expression of CD36 examined by Western blot analysis supported the suppressive effects of D-mannose against PA (Figure 6H). We have also evaluated CD36 expression *in vivo* in liver samples, and data showed that the db/db liver had increased CD36 expression compared to db/m, which was inhibited by D-mannose therapy (Figure 6I and J). These results suggest macrophage CD36 as a candidate target for mediating D-mannose effects.

**Figure 6.**
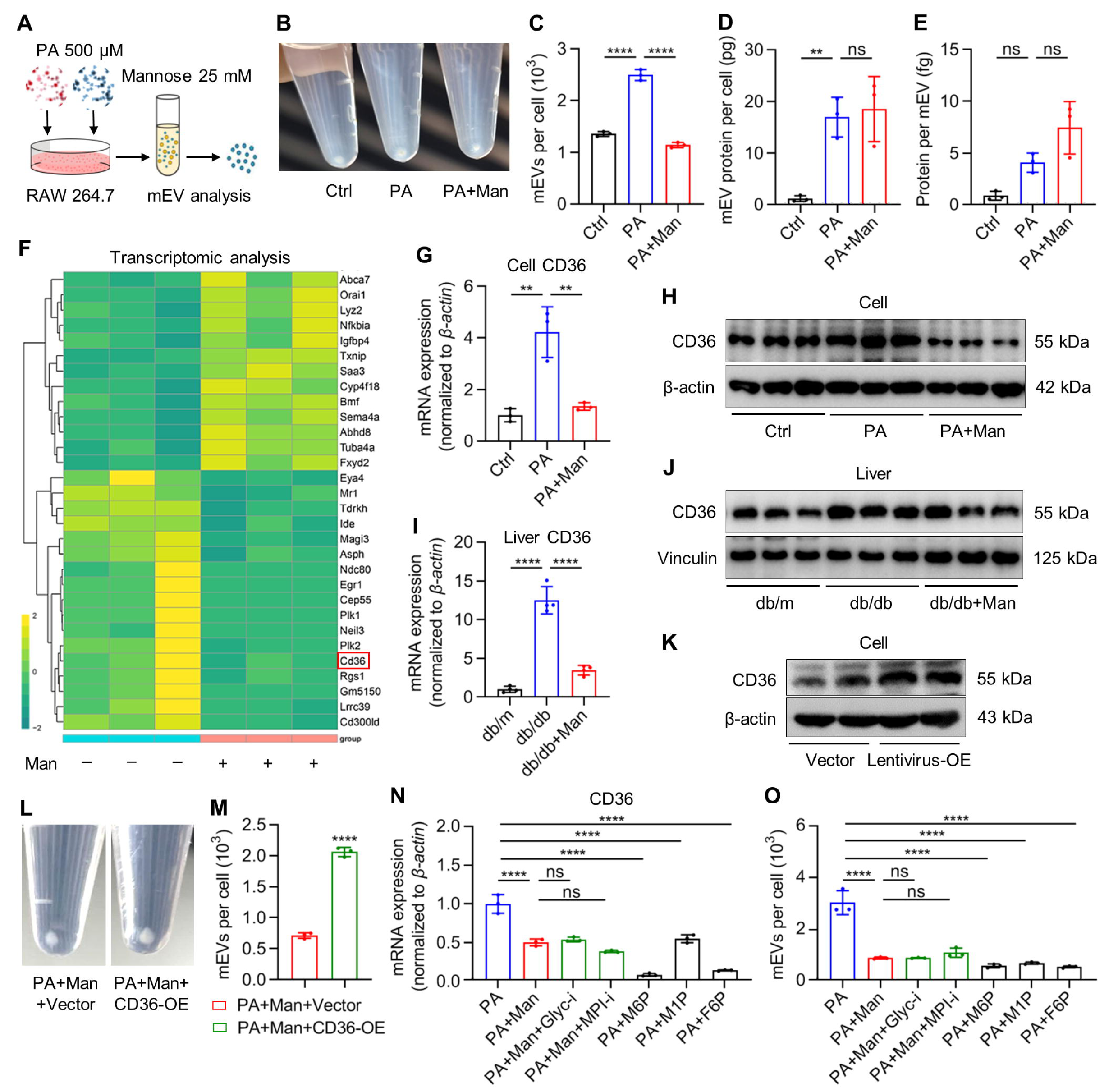
D-mannose metabolism suppresses CD36 expression to control macrophage extracellular vesicle (EV) release. (A) The diagram showing palmitic acid (PA) and mannose (Man) treatment of macrophages for analyzing macrophage-derived EVs (mEVs). (B) The gross view image of collected mEVs in 1.5-mL tubes from one 6-well macrophages. (C) Quantification of mEVs by nanoparticle tracking analysis (NTA). n=3. (D) Quantification of mEV protein content by BCA assay. n=3. (E) Quantification of protein content per mEV. n=3. (F) mRNA sequencing analysis of macrophages with or without mannose treatment. (G) Quantitative real-time polymerase chain reaction (qRT-PCR) analysis of CD36 expression in macrophages. n=3. (H) Western blot analysis of CD36 expression in macrophages. (I) qRT- PCR analysis of CD36 expression in the liver. n=4. (J) Western blot analysis of CD36 expression in the liver. (K) Western blot analysis of CD36 expression in macrophages after transfection of CD36 over expression (OE) lentivirus or its vector. (L) Gross view images of collected mEVs in 1.5-mL tubes from one 6-well macrophages. (M) Quantification of mEVs by NTA. n=3. (N) qRT-PCR analysis of CD36 expression in macrophages. n=3. Glyc-i, inhibitor of protein N-glycosylation; MPI-i, inhibitor of mannose-6-phosphate isomerase; M6P, mannose-6-phosphate; M1P, mannose-1-phosphate; F6P, fructose-6-phosphate. (O) Quantification of mEVs by NTA. n=3. Mean ± SD. **, *P* < 0.01; ****, *P* < 0.0001; ns, *P* > 0.05. Two-tailed unpaired Student’s *t* test (M) or One-way ANOVA with Turkey’s post-hoc test (C-E, G, I, N and O).

To prove that whether CD36 indeed contributed to D-mannose regulation of EV release in macrophages, we performed lentivirus-based gene over expression (OE) in macrophages, and the CD36-OE efficacy was confirmed by Western blot analysis (Figure 6K). Importantly, we revealed that CD36-OE reversed suppressive effects of D-mannose on PA-induced mEV release (Figure 6L and M), indicating that CD36 was the key to mediating D-mannose effects. To further investigate how D-mannose may regulate CD36 expression in macrophages, we applied chemical inhibitors of MPI and protein N-glycosylation to respectively block each of the two metabolic cascades of D-mannose ^16,26,27^. We have also tested metabolites of D-mannose along the cascades, mannose-6-phosphate (M6P), mannose-1-phosphate (M1P) and fructose-6-phosphate (F6P), by treating PA-conditioned macrophages instead of D-mannose. Intriguingly, we discovered that neither protein N-glycosylation nor MPI inhibition was able to counteract D-mannose to suppress CD36 expression and mEV release under PA treatment, whereas each of the M6P, M1P and F6P was enough to replicate D-mannose effects (Figure 6N and O). These above results indicate that D-mannose potently controls macrophage EV release by robust suppression of the CD36 gene expression through itself and its metabolic processes.

Taken together, the main findings of this study uncover an effective and potential T2D therapeutic by drinking-water supplementation of D-mannose, which rescued hepatocyte steatosis through suppressing macrophage release of EVs based on metabolic control of CD36 expression (Figure 7). Our findings add to the current knowledge of naturally existed sugars regulating EVs-mediated intercellular communication and shed light on translational pharmaceutical strategies of T2D.

**Figure 7.**
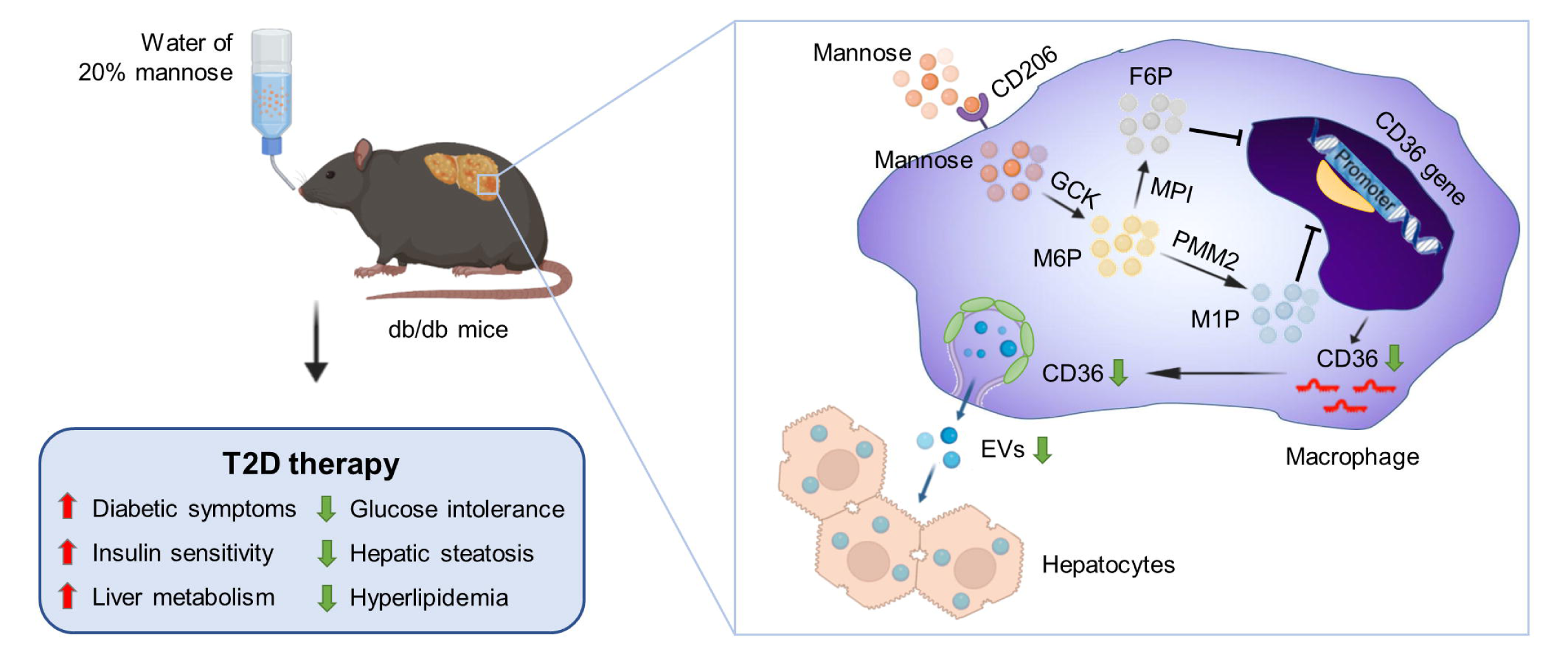
Working model of oral administration of D-mannose ameliorating type 2 diabetes (T2D) in db/db mice. D-mannose is dissolved in drinking water at 20 g per 100 mL for treating db/db mice for 8 weeks. D-mannose is internalized by macrophages *via* the CD206 receptor and is metabolized through a series of enzymes. Mannose with its metabolites is capable of suppressing CD36 gene expression, which subsequently inhibits release of hepatocyte-regulating extracellular vesicles (EVs). Accordingly, multiple symptoms of T2D are ameliorated. GCK, glucokinase; PMM2, phosphomannomutase 2; MPI, mannose- 6-phosphate isomerase; M6P, mannose-6-phosphate; M1P, mannose-1-phosphate; F6P, fructose-6-phosphate.

## 4. Discussion

The monosaccharide D-mannose exists naturally in low abundance in human blood, while an increased plasma mannose level is associated with insulin resistance and the incidence of T2D in patients ^12–14^. However, whether and how D-mannose regulates T2D development remains elusive. In this study, through a series of experiments in genetically obese db/db mice ^24^, we demonstrate that despite the altered mannose metabolism in T2D, drinking-water supplementation of supraphysiological D-mannose safely ameliorates T2D. Dysregulation of the mannose metabolism in obese individuals has been previously documented to underlie a shift in the utilization of carbohydrate substrates in the liver, but the hepatic expressions of genes responsible for mannose processing were detected with complicated results ^12^. Lee *et al*., have reported that the hepatic expression of *GCK*, the main hexokinase in the liver to converts D-mannose to M6P, was upregulated in obese subjects with co-upregulated *PMM1* and *Guanosine diphosphate (GDP)-mannose pyrophosphorylase A* (*GMPPA*)/*GMPPB*, which respectively converts M6P to M1P and M1P to GDP-mannose for the downstream N-glycan metabolism ^12^. They have also reported downregulated *Hexokinase 1* (*HK1*) and *HK2* in the liver of obese people and proposed a causal relationship between hepatic changes of these genes with the increased plasma level of mannose, hypothesizing a reduced capability of mannose consumption despite contradictory data among the detected gene expression ^12^. In the present study, we perform further analysis of the hepatic expression of genes related to mannose metabolism in db/db mice, which not only include mannose processing enzymes but also mannose synthesizing enzymes in the gluconeogenesis cascade. Our findings show upregulated expression of mannose synthesizing enzymes, *Pck1* and *Fbp1*, and confirm upregulation of *Gck* in the liver of obese subjects, which suggest increased endogenous mannose production with sustained capability of mannose consumption in T2D. Our results thus provide a progressive perspective explaining the high mannose level in T2D as only a side-effect of elevated gluconeogenesis and serve as one theoretical basis for the organism to potently utilize exogenously administered mannose for therapy.

D-mannose exists naturally in common plants and fruits, such as the broccoli, onions, cranberries and oranges, which is easily obtained and has been made ready-to-use dietary powders for food, healthcare and clinical usage ^46^. In human, long-term efficacy and safety of oral D-mannose administration have been recognized in treating MPI-COG and UTI ^17,18^. In mice, initial important works from the Chen group have revealed D-mannose as an inducer of Foxp3^+^ regulatory T cells (Treg cells) by promoting transforming growth factor-beta (TGF-β) based on increased fatty acid oxidation ^19^. Accordingly, drinking-water supplementation of D- mannose applied to improve autoimmune T1D and airway inflammation ^19^. Immunoregulatory function of D-mannose has been later documented by Torretta *et al*., to involve suppression of macrophage production of IL-1β by accumulated intracellular M6P impairing the glucose utilization, which contributed to alleviation of lipopolysaccharide (LPS)-induced endotoxemia and dextran sulfate sodium (DSS)-induced colitis in mice ^20^. The “metabolic hijack” effect of D-mannose further plays a critical role to retard tumor growth and enhance chemotherapy ^47^. Also, D-mannose has been revealed to counteract hepatic steatosis in alcoholic liver disease in mice *via* rescuing hepatocyte fatty acid oxidation and suppressing the phosphatidylinositol- 3-kinase (PI3K)/Akt/mammalian target of rapamycin (mTOR) signaling ^22^. Sharma *et al*., have additionally reported that less energy harvest by the gut microbiota partially contributes to D- mannose-mediated lean phenotype in preventing dietary obesity in mice ^21^. In this study, we surprisingly discover that providing D-mannose in early life does not ameliorate the obese phenotype in db/db mice, nor affect the gut microbiome diversity. Furthermore, D-mannose administration in our study does not influence the peripheral blood T lymphocyte percentages, nor directly target hepatocytes *in vivo*, as exhibited by fluorescent tracing images. However, drinking-water supplementation of D-mannose indeed rescues the non-alcoholic fatty liver disease (NAFLD) phenotype, improves hepatic insulin sensitivity and alleviates T2D in db/db mice, which are based on macrophage regulation but notably, independent of bioenergetic modulatory effects. Therefore, our findings add to the current mechanistic understanding of D-mannose effects, indicating gene transcription regulation by mannose and its metabolites is necessary and important. How D-mannose and its metabolites suppress CD36 expression in macrophages will require further studies.

EVs are lipid bilayered nanoparticles secreted or blebbed from almost all cell types in the body, which are loaded with a variety of signaling biomolecules, including nucleic acids and proteins ^48^. Although initially considered as cellular wastes, EVs have now been recognized as important messengers to mediate intercellular communication with emerging physiological and pathological functions ^10,11^. Notably, the metabolic homeostasis depends on the complex, multi-directional crosstalk between local and distant cells, which becomes dysregulated in metabolic diseases, such as obesity and diabetes ^49^. Accumulating evidence has supported a role of EVs in obesity-associated T2D metabolic disturbance, particularly the regional and systemic inflammation characteristics of macrophages related to adipose and hepatic stress ^50,51^. The Olefsky group has established that in obese mice, pro-inflammatory adipose tissue macrophages (ATMs) secrete miRNA-155-containing exosomes, endosome-originated EVs, to cause glucose intolerance and insulin resistance in remote organs, including the liver ^6^. They have further documented that the anti-inflammatory M2-polarized bone marrow-derived macrophages (BMDMs) secrete miRNA-690-transferring exosomes to improve target organ insulin sensitivity in HFD-induced obese mice ^52^. We have previously reported that infusion of liver macrophage-targeting EVs ameliorates the pro-inflammatory niche and rescues hepatic steatosis and T2D under dietary obesity in mice ^25^. These findings collectively suggest that macrophage-based modulation of release or composition of endogenous EVs would benefit the metabolic status despite obesity, but feasible methods are limited. In the present study, we provide a readily accessible pharmaceutical approach to control macrophage release of pathological EVs for T2D treatment, which will thus have immense translational value. Also notably, glycosylation participates in biogenesis, release and distribution of EVs, and surface D-mannose modification affects the fate and uptake of exogenously delivered exosomes ^23,53^. Moreover, well identified as a fatty acid transporter and a plasma membrane glycoprotein, CD36 is reported to regulate the ceramide formation in the caveolae to ensure release of fatty acid-containing exosome-like EVs, which is heavily modified at the post-translation level by N-linked glycosylation to mediate its trafficking to the plasma membrane and function ^45,54^. Although we focus on gene transcription regulation in this study, whether D-mannose *via* the glycosylation process influences CD36 might provide in-depth mechanistic understanding of EV release and novel therapeutic targets of T2D in future works. In summary, we uncover that drinking-water supplementation of D-mannose serves as an effective and potential T2D therapeutic, which rescued the hepatocyte steatosis through suppressing macrophage release of EVs based on metabolic control of CD36 expression. Our findings add to the current knowledge of naturally existed sugars regulating EV-based intercellular communication and shed light on novel translational pharmaceutical strategies of T2D.

## Supporting information

Figure S1. Multi-organ histological analysis reveals the safety of D-mannose therapy in db/db mice (related to Figure 2)

Figure S2. Macrophage extracellular vesicles (EVs) are characterized and regulate hepatocyte metabolism (related to Figure 5)

## Acknowledgments

This study is supported by grants from the National Natural Science Foundation of China (82301028 to Chen-Xi Zheng, 82371020 to Bing-Dong Sui and 81930025 to Yan Jin), the China Postdoctoral Science Foundation (BX20230485 to Chen-Xi Zheng) and the Young Science and Technology Rising Star Project of Shaanxi Province (2023KJXX-027 to Bing- Dong Sui). We are grateful for the assistance of the National Experimental Teaching Demonstration Center for Basic Medicine (AMFU).

