## Supplementary figures and images for "D-mannose suppresses macrophage release of extracellular vesicles and ameliorates type 2 diabetes"

### Figure S1. Multi-organ histological analysis reveals the safety of D-mannose therapy in db/db mice (related to Figure 2)

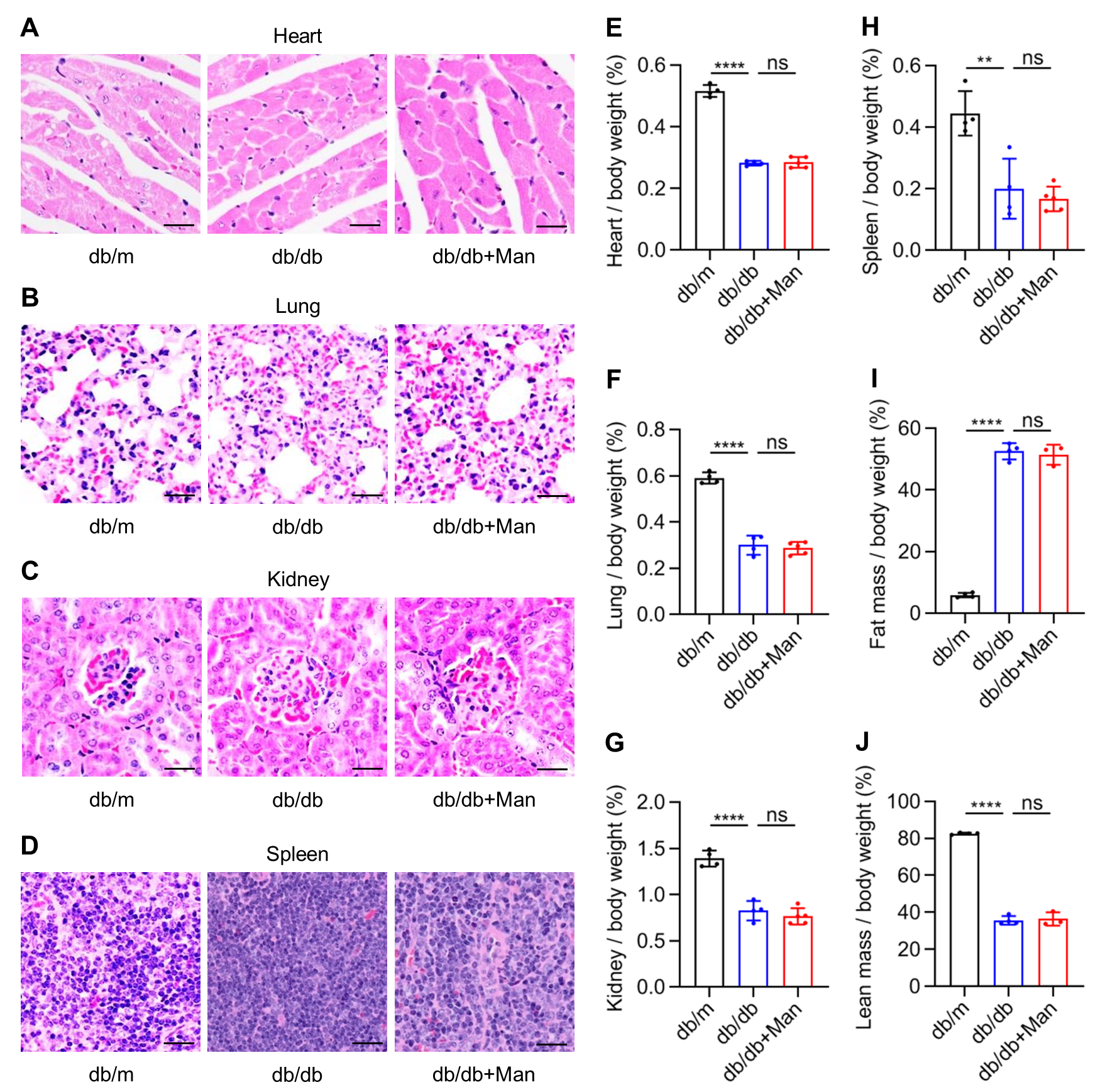

### Figure S2. Macrophage extracellular vesicles (EVs) are characterized and regulate hepatocyte metabolism (related to Figure 5)

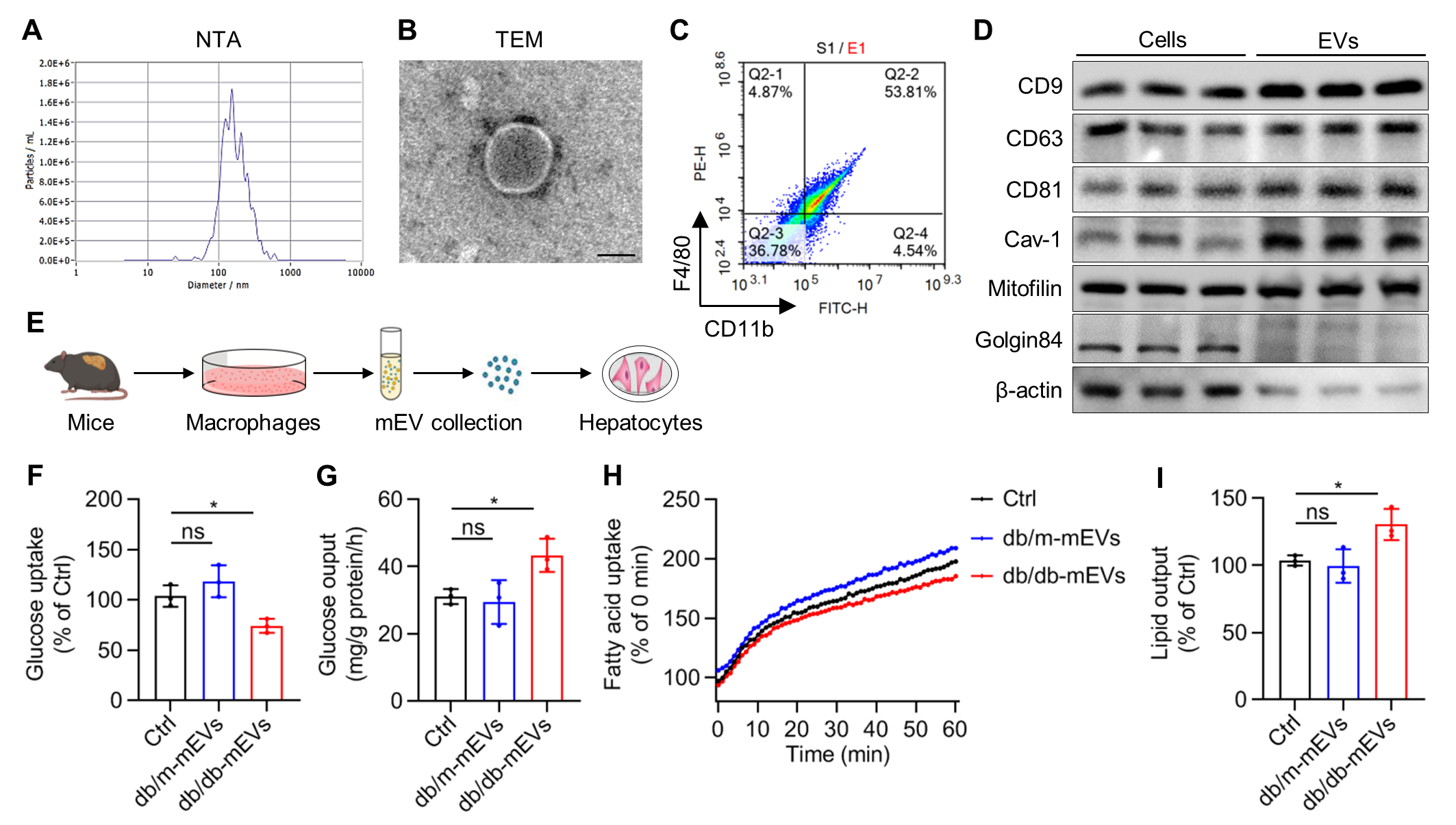
